# A simple procedure to demonstrate antimicrobial activity in cell-free supernatants

**DOI:** 10.64898/2026.06.22.733903

**Authors:** Devika Zunjarrao, Shamlan M. S. Reshamwala

**Author notes:** Corresponding author Address: Department of Biological Sciences and Biotechnology, Institute of Chemical Technology, Nathalal Parekh Marg, Matunga (East), Mumbai 400019, Maharashtra, India.

## Abstract

Probiotics produce antimicrobial peptides and small molecules that are secreted into the medium. Antimicrobial activity of cell-free supernatants can be tested using various qualitative and quantitative methods. Many of these techniques employ methods which introduce uncontrolled variables, impacting reproducibility and making comparison of reported results difficult. Here, we present a simple procedure for quantitative estimation of antimicrobial activity of cell-free supernatants which overcomes drawbacks of commonly used methods.

## Introduction

Cell-free supernatants (CFSs) of microorganisms grown in liquid media can be screened for presence of potential antimicrobial constituents using a number of well-established laboratory methods. The most commonly used methods include agar well diffusion,^1^ disk diffusion,^2^ minimum inhibitory concentration (MIC) assay^3^ and soft agar overlay bioassay.^4^ These and other methods for antimicrobial sensitivity testing form part of undergraduate microbiology laboratory courses.

Agar well diffusion method^1^ and disk diffusion method^2^ are similar techniques that are widely employed in antimicrobial susceptibility testing studies. Antimicrobial activity of a compound directly depends on its ability to diffuse through the medium, which differs from compound to compound. Since diffusion of the antimicrobial compound into the agar medium cannot be exactly ascertained, the results from these tests cannot be always reproducible and can only be a qualitative measure.^5^ Also, density of the test culture plays a role in the extent of inhibition observed.

Pre-incubation for diffusion of the antimicrobial compound into the agar may be carried out before the plate is incubated at the optimum temperature. The time and temperature of pre-incubation determine how the compound diffuses, which affects the zone of inhibition observed. A review of the scientific literature reveals that different researchers have reported using different pre-incubation times and temperatures. For example, Papagianni et al.^6^ found a three-hour pre-incubation at ambient temperature to be effective, while Malini and Savitha^7^ used an eight-hour pre-diffusion at 4°C. Choyam et al.^8^ utilized a 30-minute pre-diffusion at room temperature, Lalpuria et al.^9^ found a 48-hour long pre-diffusion at 4°C to be most effective, while Pal and Ramana^10^ used the method reported by Tagg and McGiven^11^ of pre-diffusion at 37°C for two hours. Merzoug et al.^12^ used a modification of the method reported by Tagg and McGiven^11^ and carried out a two-hour pre-diffusion at 4°C. On the other hand, many researchers, such as Goh et al.,^13^ Vimont et al.,^14^ Wu et al.,^15^ Yanagida et al.,^16^ Milette et al.^17,18^ and Abanoz and Kunduhoglu^19^ have not utilized a pre-diffusion step in their studies.

Published research papers also report using different agar well sizes. Agar well sizes of 3.5 mm,^9^ 4 mm,^10–12^ 5 mm,^6,8^ 6 mm,^17,18^ 7 mm^7,14^ and 8 mm^19^ diameter have been reported. Depth of the agar well is a variable that is mentioned only in a few studies,^9,11^ while many studies do not specify it. Volume of the antimicrobial compound added in the well also varies widely, from 25 µl,^6^ 50 µl,^8,9^ 80 µl,^12,14,17^ 100 µl^19^ to one, two and three drops.^11^

As will be clear from the above, the various parameters that affect results of the agar diffusion assay are not defined, making comparison across studies and reproducibility of results difficult.

The MIC assay,^3^ which uses liquid media and therefore avoids the pitfalls associated with the agar well diffusion assay, allows better quantification of antimicrobial activity, albeit based on visible inhibition of growth, but can often show issues with reproducibility.^20^ Variations in inoculum preparation, media, incubation temperature and time may result in variability of MIC measurement between studies.^21^ Also, results are indicated as growth (+) or no growth (-), which can be subjective as it is not quantified. Therefore, MIC may not be a strict minimum but an observable minimum which highly relies on assay parameters.^22^

The soft agar overlay bioassay is a qualitative technique first described by Gratia.^4^ It relies on proper mixing of the test microbial culture with soft agar and cooling the soft agar just enough so as to not be detrimental to the culture. It can be a bit tricky to perform and may not be suitable for organisms which do not grow robustly so as to show clear zones of inhibition.^5^ Also, antimicrobial compounds may not be produced in a quantity enough to show visible clearance. Maia et al.^22^ and Ghosh et al.^23^ describe a variation of the technique wherein spot inoculated cells are killed by exposure to chloroform before overlaying with soft agar. Incubation times following spot inoculation and soft agar overlay, respectively, are found to vary as well in the scientific literature.^22,23^

Resazurin based assays^24^ have also been utilized to test antimicrobial efficacy. However, the concentration of resazurin solution used differs between studies.^25-28^ Further, resazurin assays are limited to aerobes and microaerophiles.^29^ Moreover, resazurin has been reported to exhibit bactericidal effects towards some species.^30^

The suspension-based time-kill test standardized by the American Society for Testing and Materials (ASTM E2315-23)^31^ can be used to test biocidal potential of test materials. While very brief contact times can be studied using this method, the amount of time required for any antimicrobial substance to show visible reduction in growth of a test organism can vary from one another. The antimicrobial substance must be neutralized before assaying for viable test organisms after the contact time points. Inoculum preparation, media utilized, and contact time points selected can impact the results obtained.

To overcome the drawbacks of the widely used methods such as agar diffusion method and MIC determination, we have used a simple procedure to test antimicrobial activity of CFS of probiotic strains and compared the results with those obtained using the widely used agar diffusion method.

## Materials and Methods

### Producer and test strains

Cell-free supernatants of *Lactobacillus brevis* ATCC 367 and *Lactobacillus plantarum* MTCC 2156, as well as *Weissella paramesenteroides* D5 (GenBank accession no. OR922595) and *Enterococcus lactis* DO1 (GenBank accession no. OR922598), which were isolated from local fermented foods, were tested for antimicrobial activity against *Escherichia coli* ATCC 25922, *Pseudomonas aeruginosa* ATCC 27853 (Gram negative rods), *Listeria monocytogenes* MTCC 839 (Gram positive rod), and *Staphylococcus aureus* ATCC 25923 (Gram positive cocci).

### Media and culture conditions

Producer strains were cultivated in De-Mann, Rogosa and Sharpe (MRS) broth. Sterile Nutrient broth (NB) was used to cultivate all test organisms except *L. monocytogenes*, which was cultivated using Brain Heart Infusion (BHI) broth. Dehydrated bacterial media were from Himedia Laboratories Pvt. Ltd., India. Organisms were grown at 37°C under shaking conditions (180 rpm) for 18 hours.

### Preparation of cell-free supernatant (CFS)

18-hour old producer strain culture was centrifuged at 3900 rpm for 20 minutes at room temperature to pellet cells. Supernatant obtained after centrifugation was filter sterilized by passing through a sterile 0.2 µm Nylon syringe filter to obtain cell-free supernatant (CFS).

### Agar diffusion assay

100 µl of CFS (or MRS broth as control) was added to 7 mm diameter wells bored in Nutrient or Brain Heart Infusion (for *L. monocytogenes*) agar plates spread with the appropriate test culture. Agar was used at either 1.5% or 0.75% concentration. Plates were incubated at 37°C for 24 hours. The zone of inhibition around the wells was measured in mm.

### Determination of antimicrobial activity

Cell density of overnight cultures of the test organisms was determined using a spectrophotometer at 600 nm. Overnight cultures were diluted in 5 ml fresh sterile broth (NB or BHI) to obtain a final OD_600_ of 0.01. 1 ml CFS (or sterile broth as control) was added, and incubation was carried out at 37°C for 24 hours at 180 rpm. Growth of test organisms was determined using a spectrophotometer at 600 nm. Growth inhibition (%) by CFS was calculated by comparing cell density of the test cultures with the control (0% inhibition). All assays were carried out in triplicate with different batches of CFS.

## Results and Discussion

### Demonstrating antimicrobial activity

In this study, the effect of CFS on microbial growth was assessed. Agar diffusion, one of the most commonly used methods in the scientific literature for testing of CFS, was chosen to test antimicrobial activity of the producer strains. *L. brevis* CFS did not exhibit any activity against the organisms tested. *L. plantarum* CFS showed visible zones of inhibition against all organisms tested, while *E. lactis* CFS and *W. paramesenteroides* CFS showed activity against *E. coli* and *P. aeruginosa* only (Table 1). As the zones of inhibition observed were very small, agar concentration in the medium was decreased from 1.5% to 0.75% to allow better diffusion of the CFS into the medium. A clearer zone of inhibition was observed in the 0.75% agar plate as compared to the 1.5% agar plate but the size of the zone of inhibition remained the same (Figure 1).

**Table 1.**
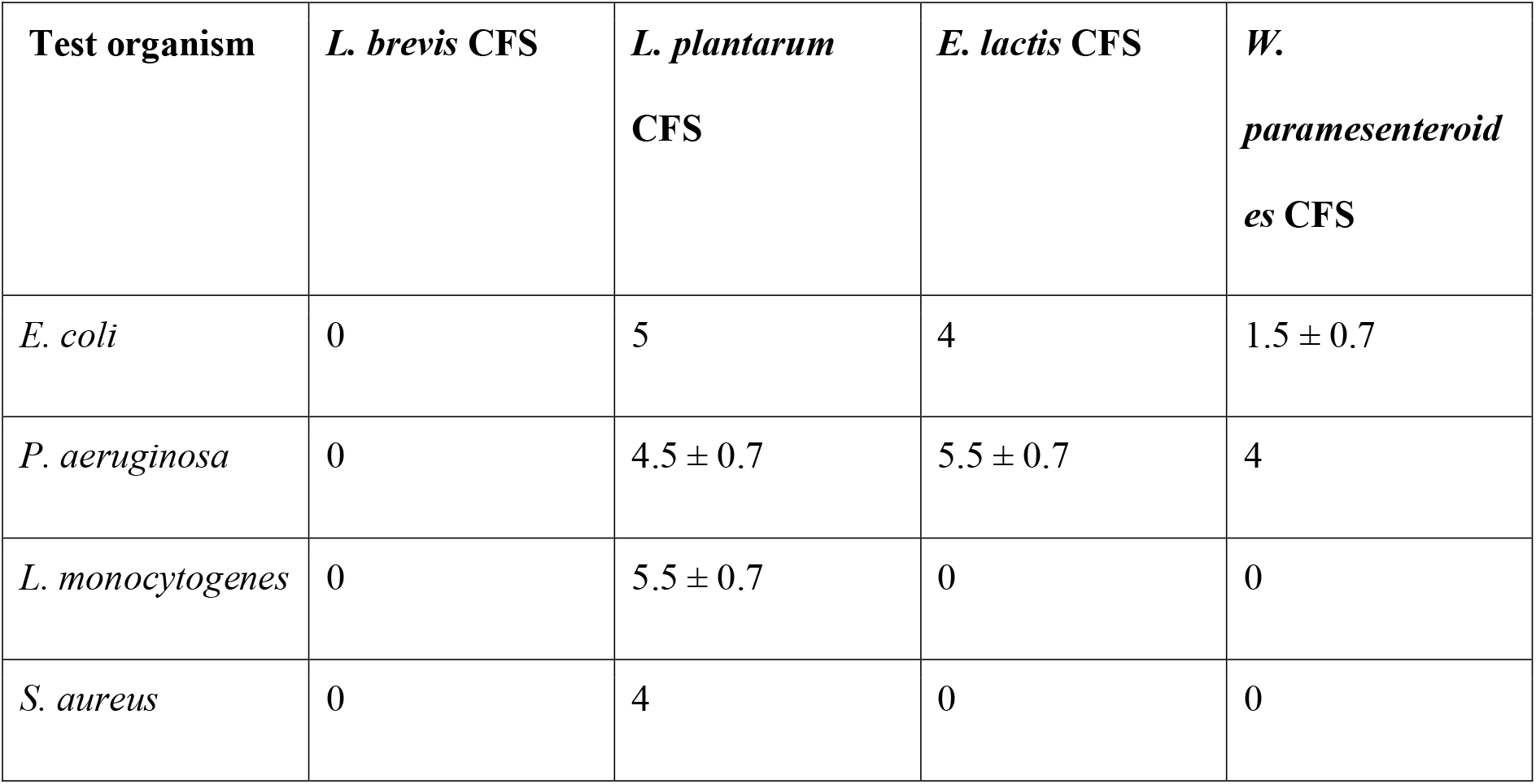
Average zones of inhibition (in mm) measured using agar diffusion method.

| Test organism | *L. brevis* CFS | *L. plantarum* CFS | *E. lactis* CFS | *W. paramesenteroides* CFS |
| --- | --- | --- | --- | --- |
| *E. coli* | 0 | 5 | 4 | 1.5 ± 0.7 |
| *P. aeruginosa* | 0 | 4.5 ± 0.7 | 5.5 ± 0.7 | 4 |
| *L. monocytogenes* | 0 | 5.5 ± 0.7 | 0 | 0 |
| *S. aureus* | 0 | 4 | 0 | 0 |

**Fig. 1.**
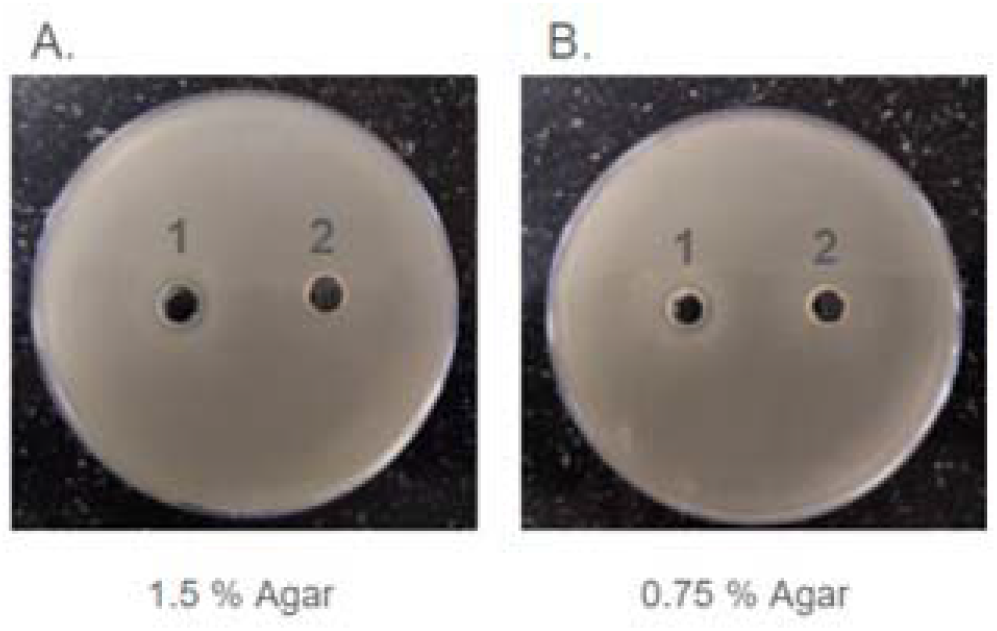
Agar diffusion assay to test antimicrobial activity of *E. lactis* CFS against *E. coli*. Well 1, *E. lactis* CFS; well 2, control (sterile MRS broth). Nutrient Agar plate containing (A) 1.5% agar and (B) 0.75% agar.

To confirm the results obtained using the agar diffusion assay, a method comprising of the following steps was used to precisely quantify growth of test organisms in the presence of CFS (see **Materials and Methods**):

1. The test organisms are inoculated in liquid medium at a cell density of OD_600_ 0.01.
2. CFS is added to the culture, and cells are incubated at their optimum growth temperature for 24 hours.
3. Cell growth is measured using a spectrophotometer, and growth inhibition (if any) is calculated based on growth observed in the control tube.

As the starting cell density is fixed and growth is measured using a spectrophotometer, reproducible results are obtained when this test is repeated.

### Antimicrobial activity testing of CFS using developed procedure

Using the method described above, it was observed that *L. brevis* CFS did not show significant inhibitory activity against the organisms tested, while *L. plantarum* and *E. lactis* CFSs showed statistically significant inhibition of all test organisms (Table 2). *W. paramesenteroides* CFS also exhibited inhibitory activity against the panel of test organisms, except *E. coli* (Table 2). It is noteworthy that the antimicrobial activity detected using our method could not be detected using the agar diffusion method, suggesting that the latter method can precisely quantify inhibition of cell growth.

**Table 2.** Average growth inhibition (%) of test organisms in presence of CFS measured using standardized assay procedure. Statistical significance of growth inhibition with respect to control is indicated

| Test organism | *L. brevis* | *L. plantarum* | *E. lactis* | *W. paramesenteroides* |
| --- | --- | --- | --- | --- |
| *E. coli* | 12 ± 0.08 % | 99 ± 0.001 % * | 98 ± 0.007 % * | 3 ± 0.03 % * |
| *P. aeruginosa* | 2 ± 0.02 % | 99 ± 0.0003 % * | 96 ± 0.01 % * | 99 ± 0.0004 % * |
| *L. monocytogenes* | 0 % | 99 ± 0.001 % * | 83 ± 0.013 % * | 98 ± 0.001 % * |
| *S. aureus* | 31 ± 0.05 % ** | 99 ± 0.0005 % * | 97 ± 0.0003 % * | 64 ± 0.021 % * |

\*P < 0.001
\*\*P < 0.01

In this study, the effect of CFS on microbial growth was assessed. However, other antimicrobial agents can also be tested using the described method. By differing the concentrations of the antimicrobial agent, the optimum concentration required to inhibit different test organisms can be determined precisely. As this assay relies on optical density measurements, even slight turbidity due to cell growth can be detected, avoiding the subjective visual assessment of growth as in the case of the MIC test.^20,21^

Additionally, the developed procedure allows quantification of extent of growth inhibition exhibited by the antimicrobial agent individually against each test organism, expressed in terms of percentage growth inhibition. This procedure for testing of antimicrobial activity overcomes the drawbacks of the widely used testing methods with respect to reproducibility and consistency of results. It provides an easy to perform, simple procedure which can be utilized for testing a variety of potential antimicrobial agents against different panels of test organisms. When testing multiple batches of antimicrobial compounds, use of the developed protocol will ensure consistent results are obtained. Results obtained using this procedure are easy to interpret and can be compared across studies to make valid deductions. Further, shelf-life or stability studies of antimicrobial agents can also be carried out by using this protocol for better comparability.

## Conclusion

In the present study, we tested antimicrobial activity of cell-free supernatants (CFS) of probiotic strains using a simple procedure that ensures reproducibility of results. The procedure is easy to perform, in which parameters such as preparation of the antimicrobial compound to be tested, test culture preparation, inoculum size, assay media volume, incubation conditions and incubation time are fixed. It overcomes the drawbacks of the commonly used methods and provides reproducible as well as quantitative estimation of antimicrobial activity.

## Statements and Declarations

### Author Contributions

SMSR conceived and designed research. DZ performed research. DZ and SMSR analyzed data. DZ wrote the manuscript. Both authors read and approved the manuscript.

## References

1. Nathan P, Law EJ, Murphy DF, MacMillan BG. A laboratory method for selection of topical antimicrobial agents to treat infected burn wounds. Burns 1978;4(3):177–177.

2. Bauer AW, Kirby WM, Sherris JC, Turck M. Antibiotic susceptibility testing by a standardized single disk method. Am J Clin Pathol. 1966;45(4):493–493.

3. Fleming A. On the antibacterial action of cultures of a Penicillium, with special reference to their use in the isolation of B. influenzæ. Br J Exp Pathol. 1929;10(3):226–226.

4. Gratia A. Numerical relationships between lysogenic bacteria and particles of bacteriophage. Ann Inst Pasteur 1936;57:652–676.

5. Balouiri M, Sadiki M, Ibnsouda SK. Methods for in vitro evaluating antimicrobial activity: a review. J Pharm Anal. 2016;6(2):71–71.

6. Papagianni M, Avramidis N, Filioussis G, Dasiou D, Ambrosiadis I. Determination of bacteriocin activity with bioassays carried out on solid and liquid substrates: assessing the factor “indicator microorganism”. Microb Cell Fact. 2006;5:30.

7. Malini M, Savitha J. Heat stable bacteriocin from Lactobacillus paracasei subsp. tolerans isolated from locally available cheese: an in vitro study. E3 J Biotechnol Pharm Res. 2012;3(2):28–28.

8. Choyam S, Lokesh D, Kempaiah BB, Kammara R. Assessing the antimicrobial activities of Ocins. Front Microbiol. 2015;6:1034.

9. Lalpuria M, Karwa V, Anantheswaran RC, Floros JD. Modified agar diffusion bioassay for better quantification of Nisaplin®. J Appl Microbiol. 2013;114(3), 663–671.

10. Pal A, Ramana KV. Purification and characterization of bacteriocin from Weissella paramesenteroides DFR-8, an isolate from cucumber (Cucumis sativus). J Food Biochem. 2010;34:932–948.

11. Tagg JR, McGiven AR. Assay system for bacteriocins. Appl Microbiol. 1971:21(5):943.

12. Merzoug M, Dalache F, Zadi Karam H, Karam NE. Isolation and preliminary characterisation of bacteriocin produced by Enterococcus faecium GHB21 isolated from Algerian paste of dates “ghars”. Ann Microbiol. 2016;66:795–805.

13. Goh HF, Philip K. Purification and characterization of bacteriocin produced by Weissella confusa A3 of dairy origin. PLoS One 2015;10(10):e0140434.

14. Vimont A, Fernandez B, Hammami R, Ababsa A, Daba H, Fliss I. Front Microbiol. 2017;8:865.

15. Wu HC, Srionnual S, Yanagida F, Chen YS. Detection and characterization of Weissellicin 110, a bacteriocin produced by Weissella cibaria. Iran J Biotechnol 2015;13(3), 63–67.

16. Yanagida F, Chen Y., Onda T, Shinohara T. Durancin L28-1A, a new bacteriocin from Enterococcus durans L28-1, isolated from soil. Lett Appl Microbiol. 2005;40(6), 430–435.

17. Millette M, Dupont C, Archambault D, Lacroix M. Partial characterization of bacteriocins produced by human Lactococcus lactis and Pediococccus acidilactici isolates. J Appl Microbiol. 2007;102(1):274282.

18. Millette M, Dupont C, Shareck F, Ruiz MT, Archambault D, Lacroix M. Purification and identification of the pediocin produced by Pediococcus acidilactici MM33, a new human intestinal strain. J Appl Microbiol. 2008;104(1):269–269.

19. Abanoz HS, Kunduhoglu B. Antimicrobial activity of a bacteriocin produced by Enterococcus faecalis KT11 against some pathogens and antibiotic-resistant bacteria. Korean J Food Sci Anim Resour. 38(5):1064–1079.

20. Kowalska-Krochmal B, Dudek-Wicher R. The minimum inhibitory concentration of antibiotics: methods, interpretation, clinical relevance. Pathogens 2021;10(2):165.

21. Mouton JW, Muller AE, Canton R, Giske CG, Kahlmeter G, Turnidge J. MIC-based dose adjustment: facts and fables. J Antimicrob Chemother. 2018;73(3):564–564.

22. Maia OB, Duarte R, Silva AM, Cara DC, Nicoli JR. Evaluation of the components of a commercial probiotic in gnotobiotic mice experimentally challenged with Salmonella enterica subsp. enterica ser. Typhimurium. Vet Microbiol. 2001;79(2), 183–189.

23. Ghosh B, Sukumar G, Ghosh AR. Purification and characterization of pediocin from probiotic Pediococcus pentosaceus GS4, MTCC 12683. Folia Microbiol. 2019;64(6):765–765.

24. Pesch KL, Simmert U. Oxidation-reduction aspects of resazurin. Milchw Forsch. 1929;8:551.

25. Sarker SD, Nahar L, Kumarasamy Y. Microtitre plate-based antibacterial assay incorporating resazurin as an indicator of cell growth, and its application in the in vitro antibacterial screening of phytochemicals. Methods 2007;42(4):321–321.

26. Foerster S, Desilvestro V, Hathaway LJ, Althaus CL, Unemo M. A new rapid resazurin-based microdilution assay for antimicrobial susceptibility testing of Neisseria gonorrhoeae. J Antimicrob Chemother. 2017;72(7):1961–1961.

27. Lescat M, Poirel L, Tinguely C, Nordmann P. A resazurin reduction-based assay for rapid detection of polymyxin resistance in Acinetobacter baumannii and Pseudomonas aeruginosa. J Clin Microbiol. 2019;57(3):e01563–18.

28. Teh CH, Nazni WA, Nurulhusna AH, Norazah A, Lee HL. Determination of antibacterial activity and minimum inhibitory concentration of larval extract of fly via resazurin-based turbidometric assay. BMC Microbiol. 2017;17(1):36.

29. Braissant O, Astasov-Frauenhoffer M, Waltimo T, Bonkat G. A review of methods to determine viability, vitality, and metabolic rates in microbiology. Front Microbiol. 2020;11:547458.

30. Schmitt DM, O’Dee DM, Cowan BN, Birch JW, Mazzella LK, Nau GJ, Horzempa J. The use of resazurin as a novel antimicrobial agent against Francisella tularensis. Front Cell Infect Microbiol. 2023;3:93.

31. ASTM E2315-23 Standard guide for assessment of antimicrobial activity using a time-kill procedure. https://www.astm.org/e2315-23.html, accessed 11 June, 2026.

